# Intraguild predation among snails drives human schistosome amplification despite susceptible host regulation

**DOI:** 10.1101/2024.12.17.629011

**Authors:** Kelsey E Shaw, Raeyan Syed, Rachel B Hartman, Danielle Mangabat, David J Civitello

## Abstract

Competitors and predators of hosts have the potential to alter transmission dynamics within host-parasite systems. Biocontrol aims to harness these effects to mitigate diseases of human or agricultural importance, but without a detailed understanding of the mechanisms of how hosts and non-hosts interact over time these attempts may backfire. We investigated the role that two resource competitors that have been employed in biocontrol, *Melanoides tuberculata* and *Physa acuta*, play in the *Biomphalaria glabrata*-*Schistosoma mansoni system*. We created experimental communities with differing compositions in mesocosms and tracked resource availability, host snail abundance, size, reproduction, and parasite production over 16 weeks. We found that *Melanoides* acted as a typical, albeit weak, resource competitor of *Biomphalaria,* reducing host body size and per capita cercarial production but with no appreciable influence on host abundance. *Physa*, however, reduced host abundance but increased resource availability and cercarial production, which represents biocontrol failure and is inconsistent with solely acting as a resource competitor. In follow-up experiments, we determined that *Physa* is a voracious consumer of host eggs. We then built a model representing this intraguild predation effect, which was able to explain these initially counterintuitive results. These divergent results from two putative resource competitors of host snails underscore the importance of establishing the mechanisms through which hosts, non-hosts, and parasites interact, especially for anticipating the potential outcomes of biocontrol interventions.

## I. Introduction

Host-parasite interactions are intricately connected to their broader ecological context. For decades, disease ecologists have focused on the role of the greater ecological community in causing either a disease “dilution effect”, in which a greater diversity of species in the environment with the host and parasite leads to decreased parasite transmission or prevalence, or the “amplification effect” in which the opposite phenomenon is observed (Civitello *et al*. 2015; Keesing *et al*. 2010; Wood *et al*. 2014). However, with increasing focus on this topic it has become apparent that knowledge of the mechanistic underpinnings of community interactions is required to predict whether dilution or amplification will occur, as either outcome is possible depending on the strength and type(s) of interaction in the host-parasite system of interest (Johnson *et al*. 2015; Rohr *et al*. 2019; Shaw & Civitello 2021). Disentangling the mechanistic relationships among hosts, parasites, and the other species they interact with is crucial for gaining a more comprehensive understanding of the community ecology of disease and the factors shaping host-parasite interactions. For example, two mechanisms have emerged that consistently drive dilution effects: transmission interference, in which nonhosts reduce host-parasite contacts, and susceptible host regulation, in which competitors of host reduce their densities (Keesing *et al*. 2006). Similarly, predators can limit disease in their prey by host regulation and by acting as culling agents, especially if they selectively attack infected individuals (Packer et al. 2003).

Here, we investigate the potential for non-host species to impact host and parasite dynamics in the snail-human schistosome system. Schistosomes are environmentally transmitted between human and snail hosts via free-living life stages that can actively seek out their host (*WHO guideline on control and elimination of human schistosomiasis* 2022). Schistosomiasis is a Neglected Tropical Disease (NTD) with devastating worldwide impacts: it is second only to malaria among parasitic NTDs in its worldwide impact on human health, and it disproportionately impacts individuals in poor, rural communities (Tucker *et al*. 2013). Despite being targeted for elimination as a public health problem by the World Health Organization 2021-2030 roadmap, decades of using conventional techniques to mitigate transmission, such as routine mass drug administration and the use of molluscicide, have not been sufficient to achieve elimination in most settings due to the lack of long-term immunity in human hosts and the potential for indiscriminate molluscicide application to lead to unanticipated feedbacks resulting in greater human transmission risk (Gryseels *et al*. 2006; Malishev & Civitello 2020; Ogongo *et al*. 2022; Wiegand *et al*. 2022). Therefore, there is increasing interest in a better understanding of ecological factors that may influence schistosome transmission and that we can potentially harness for targeted schistosomiasis mitigation in endemic regions.

The ecological community has the potential to influence snail-schistosome dynamics via multiple mechanisms. Non-host snail species have the potential to act as schistosome “decoys” that attract the parasite but then are not competent hosts, thereby acting as a sink removing parasites from the environment (Johnson *et al*. 2009). Non-host snails may also act as resource competitors. This can drive susceptible host regulation (Fenwick *et al*. 2006; Giovanelli *et al*. 2005b), but it importantly can also limit the per capita production of cercariae, the life stage of schistosomes that is infectious to people, because the parasite can be significantly limited by insufficient resources within its host (Civitello *et al*. 2022). Both of these mechanisms should reduce schistosome transmission potential, as measured by the production of cercariae by the snail population. Historically, people have attempted to exploit these mechanisms by using invasive non-host snails as biocontrol to extirpate host snails and eliminate schistosomiasis.

These efforts have resulted in local success in some regions (Pointier 1993), but in other cases have failed to control host snail populations or mitigate transmission (Lange *et al*. 2013; Pointier & Jourdane 2000; Pointier & McCullough 1989). However, most of these biocontrol efforts have occurred without long-term monitoring and data collection, making it difficult to parse which factors of either the host or the competitor snails may have resulted in this mixed success.

This study uses a mechanistic lens to investigate the impact of the ecological community on snail-schistosome dynamics and better understand why we have observed mixed outcomes in the impacts on non-host species on schistosome dynamics in natural settings. We conducted a mesocosm experiment with two putative resource competitors and schistosome decoy snail species that are invasive in schistosome-endemic settings, *Melanoides tuberculata* and *Physa acuta* (Hewitt & Willingham 2019; Laidemitt *et al*. 2022), on host snail *Biomphalaria glabrata* and *Schistosoma mansoni* dynamics. We hypothesized that both non-host species would act as resource competitors that regulated *Biomphalaria* populations, resulting in lower schistosome prevalence and decreased parasite shedding. However, tanks that contained *Physa* snails did not follow this anticipated pattern: *Biomphalaria* density decreased, but resources remained abundant and we observed sustained levels of egg laying by hosts without any appreciable recruitment of juveniles. Moreover, *Biomphalaria* coexisting with *Physa* produced significantly more human-infectious cercariae. This suite of results led to our hypothesis that *Physa* might be an intraguild predator that consumes *Biomphalaria* eggs. We then conducted egg consumption experiments that confirmed this unanticipated role of *Physa*. Last, we built a generalized model of snail-schistosome dynamics in the presence of a non-host species that could act as an intraguild predator. The model recapitulated our results that an intraguild predator can result in high levels of cercarial shedding if they are a weaker resource competitor than the host. Our results highlight the importance of understanding the underlying mechanisms at play between hosts, parasites, and species that may be acting as disease dilutors or amplifiers to predict how changing biodiversity can alter disease risk in any system.

## II. Methods

### Mesocosm set-up and sampling

We established 60L mesocosms (Rubbermaid, Wooster OH, USA) in a greenhouse in Atlanta, GA USA on March 15^th^ 2021 by filling each tank with 50L of 0.5x HHCOMBO artificial lake water and inoculating with local algae, microbes, and zooplankton, as described in (Desautels *et al*. 2022a). We supplemented 50 ug/L P, 750 ug/L N, 0.1x HHCOMBO trace solution and 2.5g dry weight of organic mushroom compost (Black Kow, Oxford, FL, USA_)_ and .5g dry weight chicken feed (Naturewise Organic Meatbird Crumbles, Nutrena, Minneapolis, MN, USA) as a carbon source weekly throughout the experiment. We also added deionized water weekly to compensate for evaporation and return the tanks to 50L; this volume was typically less than 5L. On April 1^st^ we homogenized algal communities by pooling 100mL from each tank and redistributing it.

Snails were introduced into tanks on April 15^th^ 2021 and sampled weekly until August 10^th^ 2021 (117 days). April 15^th^ is considered day 0 in week 0 of the experiment, and sampling concluded after week 16. In week 0, 10 adult *Biomphalaria glabrata* snails (host of *Schistosoma mansoni*) between 9-12mm shell length were added to all tanks, and four treatment groups were formed in an additive design as follows: i) *Biomphalaria* only, 10 founder snails ii) *Biomphalaria* plus 10 size-match *Melanoides tuberculata* iii) *Biomphalaria* plus 10 size-match *Physa acuta* iv) *Biomphalaria* plus 10 *Melanoides* and 10 *Physa* snails. *Melanoides* and *Physa* snails were chosen because they are not competent hosts for *Schistosoma mansoni*, but are putative resource competitors with *Biomphalaria* and potential schistosome decoy species, and these genera co-exist in schistosome endemic regions (Hewitt & Willingham 2019; Laidemitt *et al*. 2022). Initial snail densities were chosen to represent small founding populations observed in natural populations and based on mesocosm dynamics we have observed in previous experiments (Desautels *et al*. 2022a; Wood *et al*. 2019). *Biomphalaria* were originally obtained from the Schistosomiasis Resource Center, Rockville MD USA; *Physa* were obtained from the Atlanta Water Gardens, Atlanta GA USA; and *Melanodies* were obtained from Aquatic Discounts, Ej Cajon CA, USA.

We obtained mice infected with NMRI-strain *S. mansoni* from the Schistosomiasis Resource Center and used 5 mouse livers per parasite inoculation. We introduced *S. mansoni* eggs into each tank on days 5, 27, 48 and 69. Eggs were purified following the protocol from (AU - Dinguirard *et al*. 2018) and distributed into 6mL aliquots of volume per tank. We sampled 4 extra aliquots per dose to estimate egg abundances which were: 2520 ± 373.10, 1830 ± 164.23, 1335 ± 149.75, and 3795 ±1 53.70 (mean ± s.e.) eggs per tank.

On day 11 (week 1), we began weekly sampling for snail abundance, body size, and reproduction. Using a handheld net, we exhaustively censused each tank, then passed what we collected through a 2mm mesh to exclude neonatal snails. Snails were photographed and returned to their tanks, and abundance and body size were later recorded using Fiji image analysis software (Schindelin *et al*. 2012), using a fixed grid in the photo for calibration. Snail reproduction was estimated by counting the number of eggs for *Biomphalaria* and *Physa* (easily visually distinguishable) laid on two 10cm x 20cm plexiglass tiles submerged in each tank; all eggs were scraped off each week before replacing the tile in the water. *Melanoides* exhibit live birth, and thus we were unable to estimate their reproduction as a separate quantity from abundance. Starting on day 18, we measured photosynthetic periphyton growth every two weeks as a proxy for resource availability to the snails via *in vivo* flourescence as described in (Desautels *et al*. 2022a).

Cercariae, the life stage of schistosomes which is infectious to humans, were measured beginning on day 32, 4 weeks after the first introduction of schistosome eggs (the typical minimum prepatent period in the snail host under good conditions). All *Biomphalaria* collected for each tank were placed in a 500mL Erlenmeyer flask with 400mL of 0.5x HHCOMBO and set in the sunlight for 90 minutes, after which time snails were returned to their tanks. All cercarial sampling occurred between 9 and 11 am to minimize the impact of circadian emergence confounding our results (Wolmarans *et al*. 2002). Once snails had been removed from the flask, the remaining solution was passed through a 10um filter using vacuum filtration to concentrate the cercariae. They were then resuspended in 25mL of 0.5x HHCOMBO and immediately preserved and stained with 2mL of Lugol’s iodine, and later counted under a dissecting microscope (40X). However, we did not detect many infected snails and thus have omitted tank-level parasite data from our analyses. Instead, we analyzed *per capita* cercarial release of uniquely marked sentinel snails.

### Sentinel snail set-up and sampling

In weeks 4-8 very little cercarial production was observed in the tanks, regardless of snail community makeup, and we became concerned that poor infection success for any number of reasons would compromise the experiment. In response, we initiated controlled infections of individual “sentinel” snails to allow us to observe the impact of snail community on parasite production, on day 63 (week 9) we introduced 8 sentinel *Biomphalaria* snails per tank. Sentinels were between 6 and 8 mm in shell length at the time of introduction and painted with a unique color to allow for individual-level data collection in this group in a mark-recapture design.

Sentinels were exposed to 8 freshly hatched schistosome miracidia in 5 mL of 0.5x HHCOMBO the day before introduction into the mesocosm. For this size of snail and number of miracidia, based on previous experiments in our lab, we anticipate between a 60-85% prevalence of infection (Shaw *et al*. 2022). In addition to sentinel snails added to the mesocosms, 10 additional sentinel snails were maintained in lab on high quality *ad libitum* food consisting of 4 g homogenized chicken feed (Naturewise Meatbird Crumbles, Nutrena Animal Feeds, Minneapolis, Minnesota, USA) and 2 g fish flakes (Omega One Freshwater Flakes, OmegaSea, Painesville, Ohio, USA) embedded in 1 g agar dissolved in 90 mL deionized water (Desautels *et al*. 2022b).

After introduction to the tanks, sentinel snails were sampled weekly as part of the exhaustive tank censuses and their individual body size as well as survival status was recorded. Beginning in week 13, sentinel snail cercarial production was measured by placing sentinel snails individually in 20mL of 0.5x HHCOMBO and placing them in a bright light for 90 minutes. Sentinels were then replaced in the tanks and the cercariae were immediately preserved and stained with 2mL of Lugol’s iodine, and later counted under a dissecting microscope (40X).

### Analysis of population and community dynamics

We analyzed periphyton production, *Biomphalaria* egg production and *Biomphalaria* abundance with Generalized Additive Mixed Models. GAMMs are a powerful tool to investigate potentially non-linear dynamics of the different snail communities over time (Desautels *et al*. 2023). For each response variable, we fit a model that predicted the time course as a smooth function with the *Biomphalaria*-only treatment as a baseline, s(Time), plus an additional smooth function of time for the difference between the other snail communities (“Treatment”) and the *Biomphalaria*-only populations, resulting in the model syntax (Equation 1):

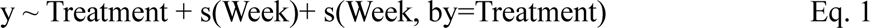

Thus, the significance of any of these “difference smooths” indicates divergent temporal dynamics between that treatment and the *Biomphalaria*-only control.

To assess the impact of snail community on *Biomphalaria* population size structure, we calculated the cumulative distribution function of *Biomphalaria* body size in each tank, each week and then calculated the area under the curve of this distribution using the trapezoid method and the auc function in the “pROC” package (Makowski *et al*. 2019). This quantity represents the relative size structure independent of abundance. It is small when the population is dominated by small individuals and large when the population is dominated by large individuals. We then fit a GAMM with these areas under the curve as the response variable, with the same model syntax as Equation 1.

All GAMMs included a random effect of tank and the continuous autoregressive-1 error structure to account for repeated measurements of the tanks. GAMMs for snail abundance and eggs used the quasipoisson error distribution with log link to account for potential overdispersion in count data. The GAMM for periphyton production and size structure distribution used the Gamma error distribution with log link to account for potential overdispersion in continuous data. We visualized the model-predicted means ±1 SE for each treatment using predict.gam (R package mgcv, (Wood 2017) and plot_smooths (R package itsadug, (Van Rij *et al*. 2015).

### Analysis of sentinel snail dynamics

To assess the impact of snail community on sentinel snail growth over the time course of the experiment, we used GAMMs with the response variables of sentinel shell length and a nested random effect of snail within tank. GAMMs were conducted as described in the tank-level analysis, with a Gamma error distribution and log link to account for potential overdispersion in continuous data. We tested for differences in weekly cercarial production over the course of the experiment with a GAMM as in Eq. 1 with a quasipoisson error distribution and a nested random effect of snail within tank. To test for differences in sentinel snail survival, we used the Cox proportional-hazards model (pack-age: KMsurv, function: coxph) with treatment group as a predictor and tank as a random effect (Klein & Moeschberger 2003).

### Egg predation experiments

After the conclusion of the mesocosm experiment, we conducted a laboratory experiment to investigate the potential for *Biomphalaria*, *Melanoides* and *Physa* to eat *Biomphalaria* eggs. Large (20-30mm), *ad libitum* fed *Biomphalaria* were placed in 25mL beakers of 0.5x HHCOMBO for 3 days and then removed. The number of eggs laid were then counted, and cups were subjected to one of the following treatments: (i) remaining in snail-conditioned HHCOMBO with no snail to test for the potential impact of snail-associated microorganisms on eggs (ii) addition of one 8-10 mm *Biomphalaria*, (iii) addition of one 8-10 mm *Melanoides* or (iv) addition of one 8-10 mm *Physa*. All beakers were maintained at 26° C for three more days, at which time snails were removed and the number of unhatched eggs and neonate Biomphalaria were counted. The difference between the initial egg count and the number of eggs/neonates observed remaining was presumed to be predated. This set-up was repeated across three blocks. To compare the number of eggs eaten among the species, we used the glmmTMB package in R (Brooks *et al*. 2017) to fit a generalized linear mixed model (GLMM) accounting for the random effects between individual snails and block dates (bionomial probability distribution with a logit-link function). We then used the emmeans package to conduct an estimated marginal means (least-squares means) *post hoc* test (Lenth 2022).

### A model of intraguild predation

To explore the range of effects a competitor/intraguild predator (hereafter IGP), non-host snail species may have on host snail and schistosome dynamics, we built a model of a hypothetical IGP based on *Physa* interacting with *Biomphalaria* and *S. mansoni*. The model tracks the density of resources, *R*, host snail eggs, *G*, susceptible snail hosts, *S*, exposed host snails, *E*, infected snail hosts, *I*, non-host IGP snails, *IGP*, and schistosomes in the life stage infectious to snails, *M*, and the cercarial lifestage which is infectious to humans, *C*.

Resources grow logistically with a carrying capacity of *K* and growth rate *r*. Resources are consumed by host snails at rate *f_h_* and IGP snails at rate *f_IGP_*. Host snail eggs are produced based on the resource consumption of susceptible, exposed, and infected hosts with conversion efficiency ρι and infected hosts have partial castration mediated by parameter ρ, which represents the relative fecundity of infected hosts. Eggs decrease by a background death rate, *d_g_*, hatching at rate μ, and from predation by IGP snails at rate α. Susceptible hosts increase from eggs hatching at rate μ and decrease from background death (*d_s_*). Susceptible hosts move into the exposed class based on the density of miracidia, *M*, and the transmission rate which is a product of host exposure, ε_h_, and host susceptibility, σ. Exposed hosts may die from background mortality or move into the infected class at rate *l*. Infected hosts may die from background mortality or due to parasite virulence, ϖ. IGP snails increase based on resource competition with conversion efficiency ρι and consumption rate *f_IGP_* on periphyton and α on host eggs. IGP snails die at background rate *d_IGP_*. Miracidia are introduced into the environment at a constant rate, λ, and either die at background rate *m* or encounter a host snail at rate ε_h_ or a IGP snail at rate ε;_IGP_. Non- host snail genera, including *Physa*, have been shown to be effective decoy species for schistosomes and this is accounted for by the ε_IGP_ parameter (Laidemitt *et al*. 2022). Miracidia that enter IGP snails are removed from the environment but do not create successful infections.

Finally, infectious cercariae are released into the environment by infected hosts based on resource density, represented by the linear function 8\**R* (Civitello *et al*. 2018), and die at rate *m*. We ran a simulation of this model across a range of values for α and for 120 days, equivalent to the length of our mesocosm experiment. Parameter values are described in Table 1.

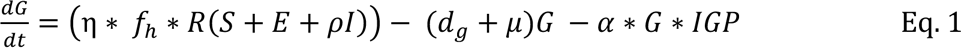

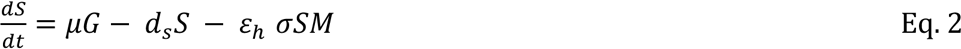

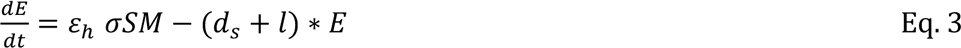

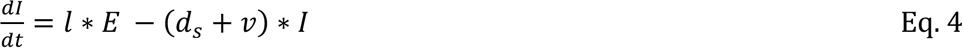

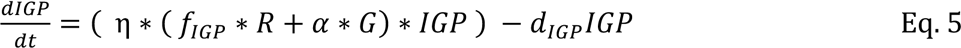

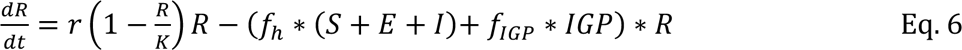

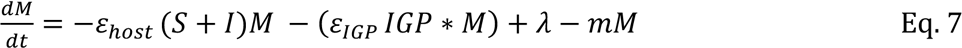

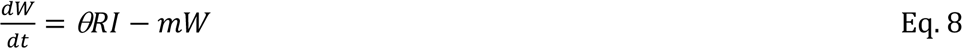

**Table 1.** Parameter values and descriptions.

| Parameter | Description | Value | Source |
| --- | --- | --- | --- |
| $\eta$ | Conversion efficiency | 0.08 | (Civitello *et al.* 2018) |
| $f_h$ | Feeding rate of host | 0.2 | (Civitello *et al.* 2018) |
| $\rho$ | Partial castration due to infection | 0.9 | (Civitello *et al.* 2018) |
| $\mu$ | Egg hatching rate | 0.071 | Personal observations of 14-day hatching period |
| $d_g$ | Death rate of host eggs | 0.036 | Personal observations of ~67% hatching rate |
| $ds$ | Death rate of host snail | 0.006 | (Civitello *et al.* 2018) |
| $\varepsilon_h$ | Exposure rate of hosts to parasites | 0.01 | (Shaw *et al.* 2022) |
| $\sigma$ | Host susceptibility to infection | 0.4 | (Shaw *et al.* 2022) |
| $l$ | Rate of moving from Exposed to Infected class | 0.045 | (Malishev & Civitello 2019) |
| $\alpha$ | Rate of predation by competitor/predator | Range from 0-0.08 | varied |
| $v$ | Parasite virulence | 0.04 | (Civitello *et al.* 2018) |
| $f_{IGP}$ | Feeding rate of competitor/predator | 0.09 | (Saha *et al.* 2017) |
| $d_{IGP}$ | Death rate of competitor/predator | 0.02 | (Saha *et al.* 2017) |
| $r$ | Growth rate of resources | 0.5 | assumed |
| $K$ | Carrying capacity of resources | 10 | assumed |
| $\epsilon_{IGP}$ | Exposure rate of competitor/predator to parasites | 0.008 | assumed |
| $\lambda$ | Rate that miracidia enter the environment | 2 | assumed |
| $m$ | Background death rate of parasites (miracidia and cercariae) | 1 | (Civitello *et al.* 2018) |
| $\theta$ | Modification parameter for the impact of resource availability on cercarial production | 100 | (Civitello *et al.* 2018) |

To explore the effect of different IGP snail parameter combinations on schistosome transmission risk, we calculated both the total cercariae produced over the course of the simulation and the R* value (Eq. 9) for the IGP snail from all possible combinations of α and *f_IGP,_* and α and *d_IGP_* parameter values over a grid of reasonable values (α = [0, 2.5], *f_IGP_*= [0, 0.4] *d_IGP_* = [0.0025, 0.025]).

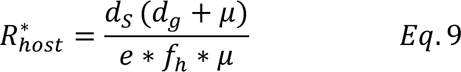

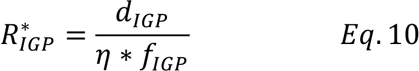

## III. Results

### Tank-level analysis

In the *Biomphalaria*-only treatment, periphyton, a proxy for snail resource availability, peaked in Week 12 of the (Time smooth, p=0.00409, Fig. 1a). Tanks with *Physa* and *Biomphalaria* had greater amounts of Periphyton over the course of the experiment (*Physa* difference smooth, p=0.0131).

**Figure 1.**
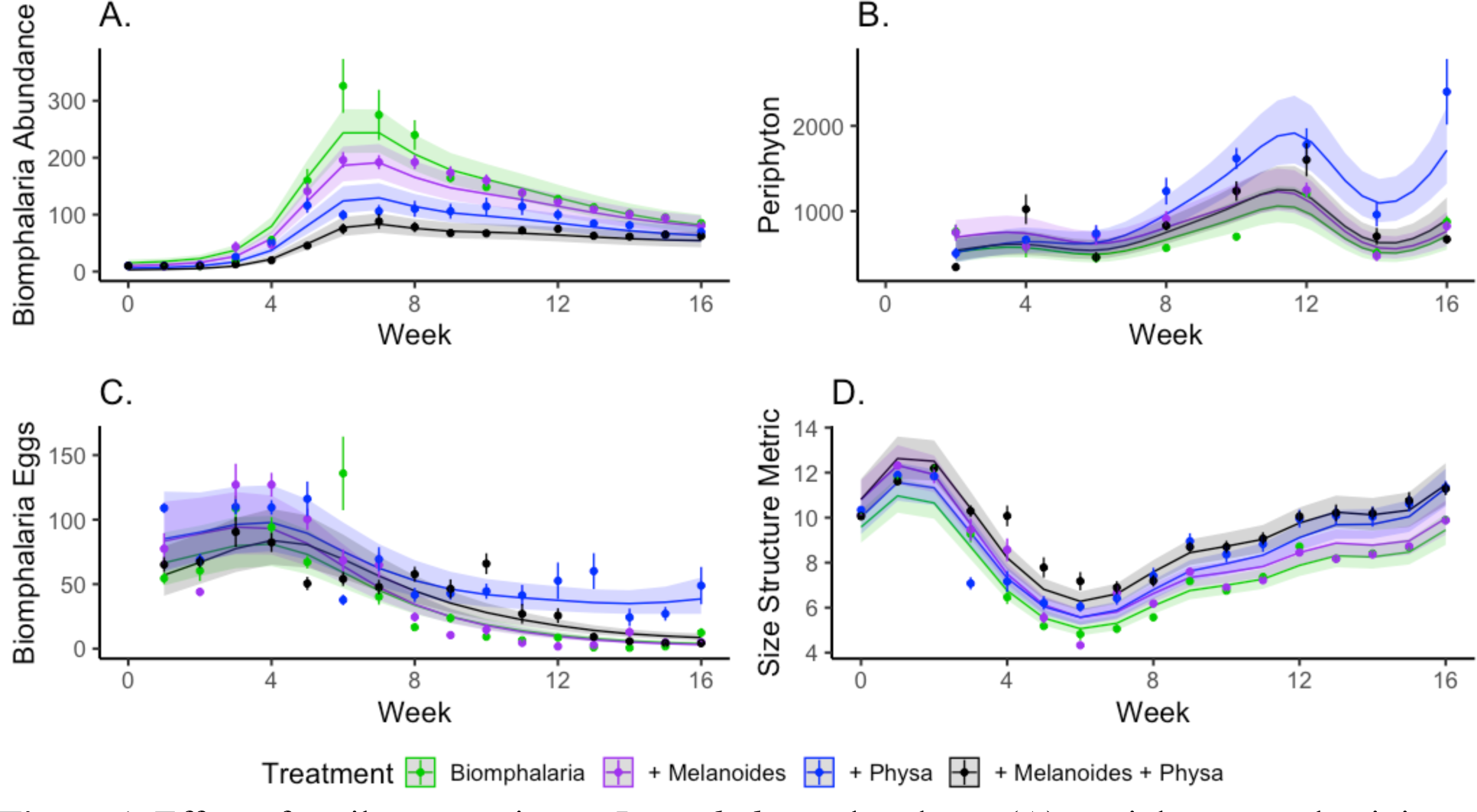
Effect of snail community on *Biomphalaria* abundance (**A**), periphyton productivity (**B**), *Biomphalaria* egg production (**C**), and *Biomphalaria* population size structure (**D**). Points and error bars are treatment means ± standard error (SE). Lines and shaded regions are model fits ± SE. (**A**) *Biomphalaria* abundance peaked and then decreased over the course of the experiment, with lower abundances in tanks that contained *Physa* as well as tanks with both *Physa* and *Melanoides*. There was significantly higher periphyton production (**B**) and higher *Biomphalaria* egg production (**C**) in tanks that contained *Physa.* (**D**) Tanks with *Physa* and *Melanoides* also had size-structures (AUC) shifted towards larger individuals.

*Biomphalaria* egg production was high in the *Biomphalaria*-only treatment over the first six weeks of the experiment before declining (Time smooth, p<0.001, Fig. 1b). Relative to *Biomphalaria*-only tanks, egg production was highest in tanks that contained *Physa* and *Biomphalaria* snails (*Physa* difference smooth, p = 0.0028), and remained higher over time (*Physa* difference smooth, p<0.001).

*Biomphalaria* abundance peaked in *Biomphalaria*-only tanks in Weeks 6-8 before declining (Time smooth, p<0.001, Fig. 1c), and relative to tanks that only contained *Biomphalaria*, tanks with *Biomphalaria*, *Physa*, and *Melanoides* had lower *Biomphalaria* abundances throughout the experiment (Treatment smooth, p=0.0187). Tanks that contained *Biomphalaria* and *Physa* (t=- 2.768, p=0.00584) or *Biomphalaria*, *Physa*, and *Melanoides* (t=-4.204, p<0.001) had significantly lower abundances of *Biomphalaria* compared to *Biomphalaria*-only tanks. Interestingly, tanks with *Biomphalaria* and *Melanoides* were not significantly different from tanks with *Biomphalaria* alone (t=-1.154, p=0.249).

Population size structure shifted from medium-sized founder snails to being dominated by small individuals (smaller Area Under the Curve) in Weeks 4-8 of the experiment, then over time returned to populations dominated by larger individuals (Time smooth, p<0.001, Fig. 1d). Tanks with *Biomphalaria*, *Physa*, and *Melanoides* snails had significantly higher Areas Under the Curve, indicating a population with a greater proportion of large *Biomphalaria* compared to *Biomphalaria*-only tanks (t=2.578, p=0.0102).

### Individual-level analysis

Shell length in all groups generally increased over time (Time smooth, p<0.01, Treatment smooth for *Melanoides*, p=0.0275, for *Physa* p<0.001, for *Melanoides* and *Physa* combined p<0.001) (Fig 2a). Sentinel *Biomphalaria* shell length was significantly smaller in tanks with *Melanoides* snails (t=-2.001, p = 0.0456). Cercarial output in all groups decreased over time (Time smooth, p<0.0001, Treatment smooth for *Melanoides*, p=0.000169, for *Physa* p=0.0146, for *Melanoides* and *Physa* combined p=0.0206). Sentinel snails in tanks that contained Melanoides shed significantly fewer cercariae (t=-2.233, p=0.0262), while those in tanks with Physa shed significantly more cercariae (t =4.212, p=0.0000321; Fig 2c). Sentinel snails in tanks with both *Physa* and *Melanoides* had the lowest mortality throughout the experiment (z=-2.693, p=0.00707; Fig 2d).

**Figure 2.**
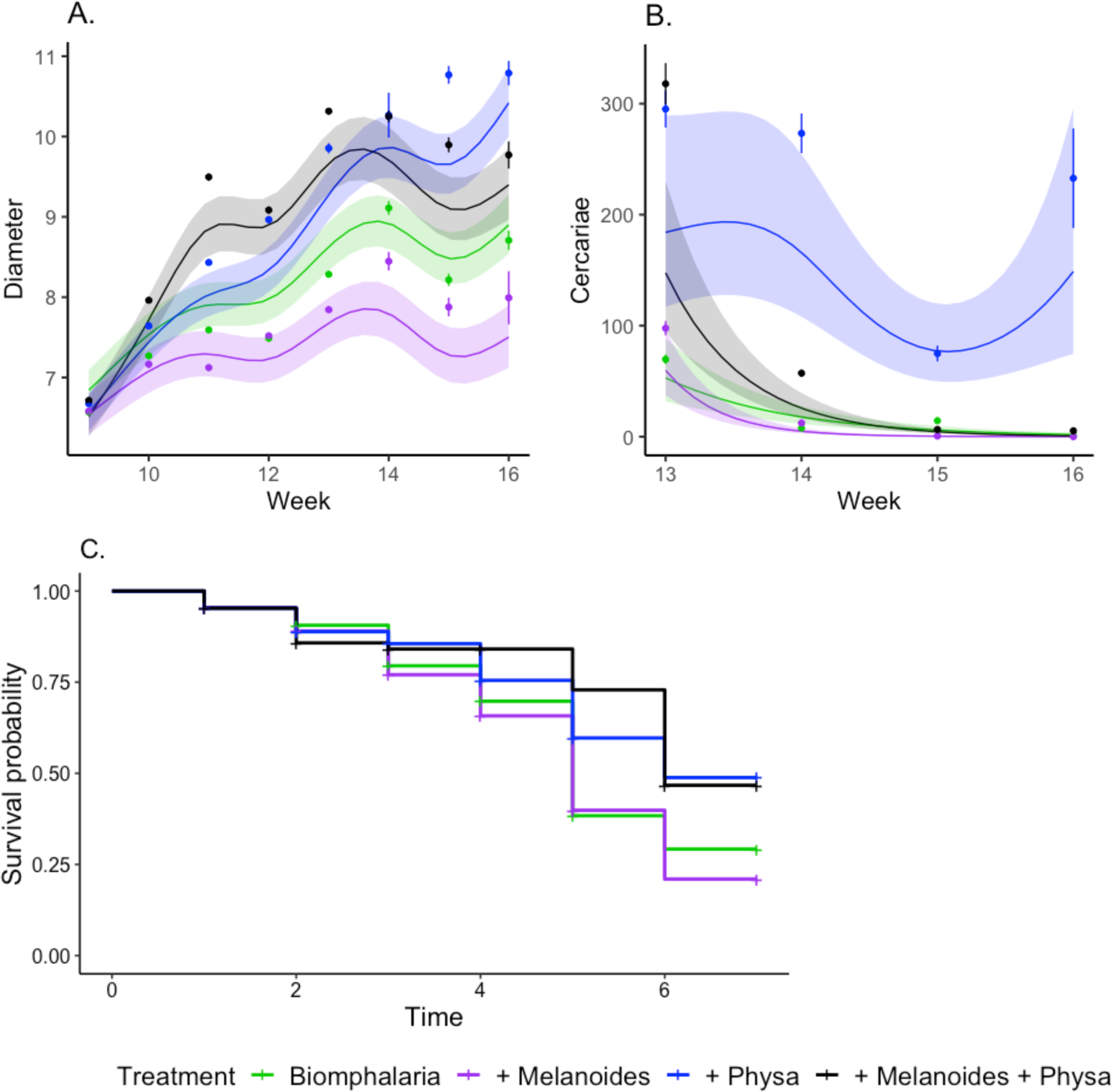
Sentinel *Biomphalaria* shell length (**A**) generally increased over time, but was smallest in tanks that contained *Melanoides*. (**B**) Sentinel snails in the tanks with *Melanoides* shed significantly fewer cercariae than tanks with *Biomphalaria* alone, while tanks with *Physa* shed significantly more. (**C**) Sentinel snails in tanks with both *Physa* and *Melanoides* had lower mortality compared to *Biomphalaria*-only tanks.

### Egg predation

We found that *Physa* snails were avid predators of *Biomphalaria* eggs (Fig. 3). Cups with *Physa* had significantly more egg destruction than snail conditioned water (z value = 8.397, p<0.0001). *Melanoides* and *Biomphalaria* cups did not significantly differ from snail-conditioned water (*Melanoides*: z value = 0.278, p=0.781; *Biomphalaria*: z value = 0.387, p=0.699). Pairwise post- hoc testing revealed that *Physa* also ate significantly more eggs than *Biomphalaria* (z ratio = - 4.649, p <0.001) and *Melanoides* (z ratio = -4.303, p= 0.001).

**Figure 3.**
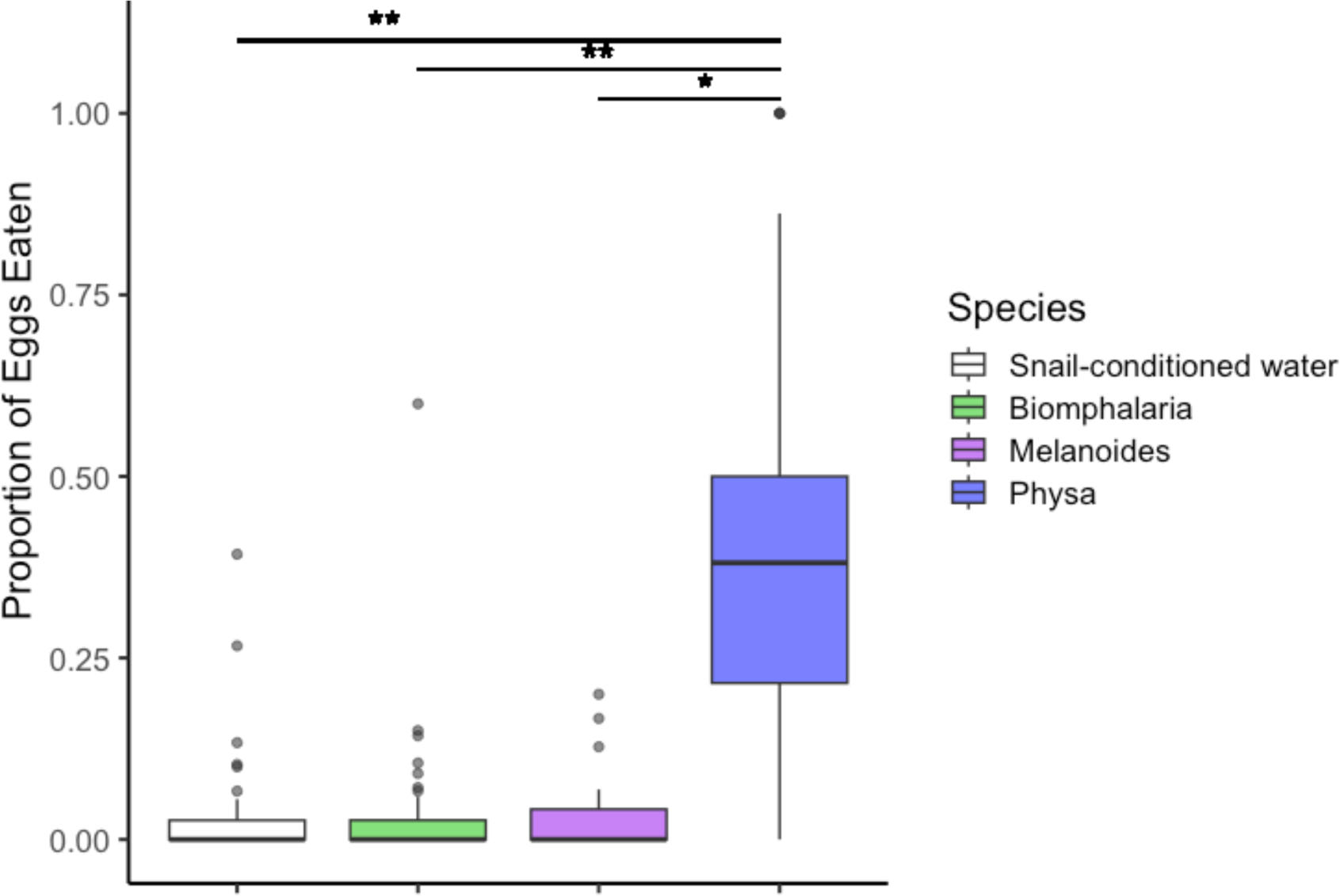
*Physa* snails were voracious predators of *Biomphalaria* eggs, consuming significantly more eggs than another other group (n=25 *Physa*, 23 *Melanoides*, 25 *Biomphalaria* and 15 snail- conditioned control). * represents p=0.0001, ** is p<0.0001

### A model of intraguild predation

We found that, in the absence of predation, our model captures classic dilution effects when the feeding rate of the competitor is greater than that of the host: susceptible host density is regulated and cercarial production goes down both due to lower host density and lower resource availability (Fig 4A). Once we included predation in the model, we found that intraguild predator (IGP) density at the end of the experiment still exceeds that of both susceptible and infected snails similar to the no-predation scenario (Fig 4B). However, despite the smaller host snail density, the partial release of host snails from intraspecific competition results in greater resource availability and therefore a higher production of infectious cercariae for a range of α values (Fig 4B). Therefore, there is a range of values for α that result in the presence of an intraguild predator increasing human hazard of encountering schistosomes, similar to what we found in our experimental mesocosms.

**Figure 4.**
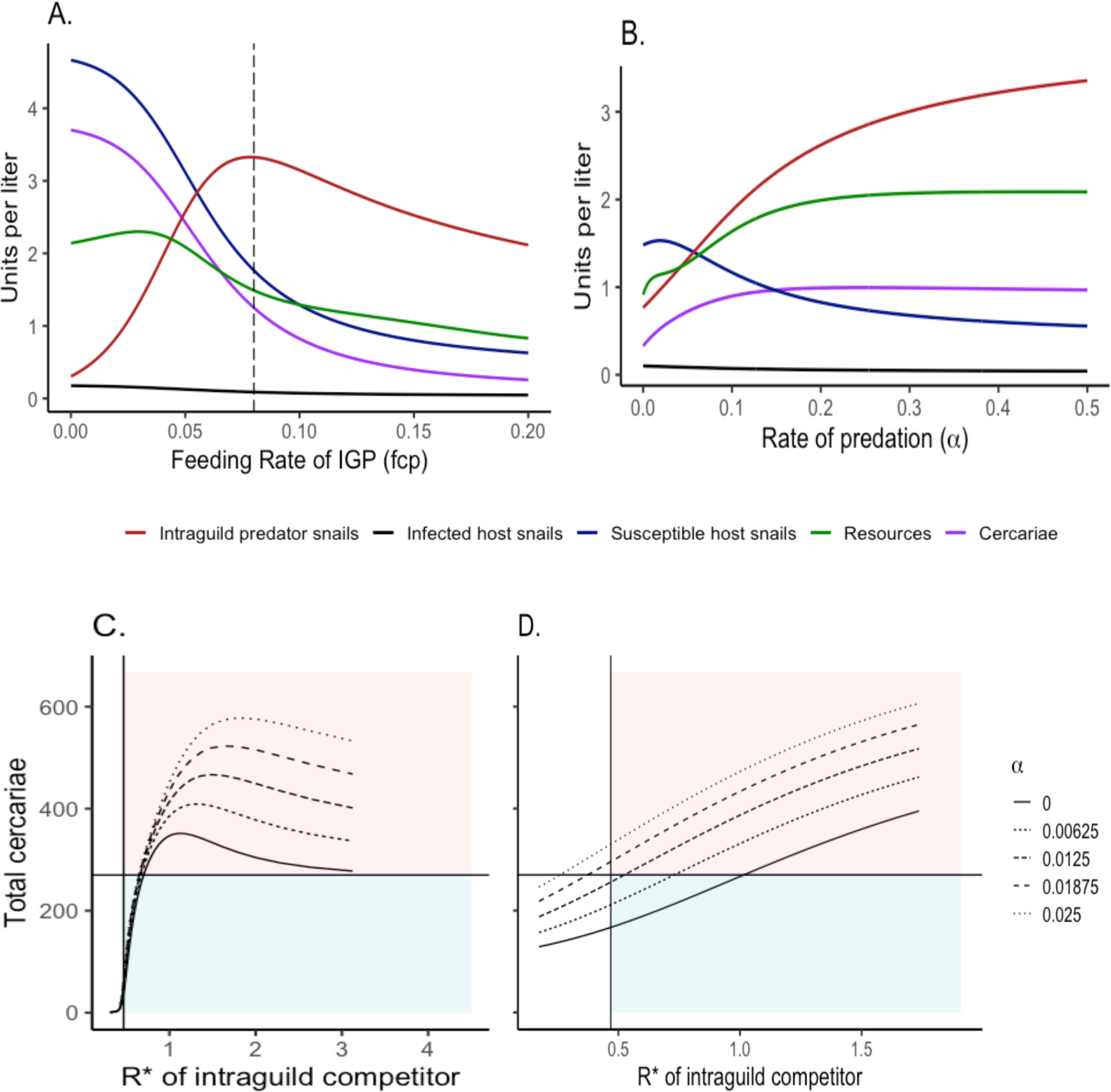
**(A)** Model outputs in the absence of predation (α=0) and variable competitor feeding rates (*f_cp_*) of the equilibrium density of susceptible and infected host snails as well as the equilibrium density of IGP snails (individuals/liter), resources (mg of algae per liter), and the summed cercarial output from the entire simulation (individuals/liter, scaled by a factor of 1/1000 to fit to the same scale). Vertical line represents host feeding rate (*f_h_* = 0.08). As competitor feeding rate increases, host snail densities decrease along with cercarial production. (**B)** Same model outputs as in (A), but with a varied rate of predation (α) and fixed competitor feeding rate (*f_cp_ =* 0.08). Above a predation rate of approximately 0.08, despite intraguild predator snails outcompeting host snails, cercarial output remains high due to a greater availability of resources. (**C)** Simulation of total cercarial production by host snails across various R* values of the IGP based on varying α and *f_cp_* ; points represent output of simulations. Vertical line represents the R* of the host snail and the horizontal line represents the total cercariae that hosts would produce in the absence of the IGP. Dilution (blue shading) is observed in the presence of the IGP in a small set of simulations in which the R* value of the IGP is low; however, for a majority of the simulations across parameter values, amplification (red shading) is observed in which the R* value of the IGP is greater than that of the host and there is a greater cercarial output then would be expected in the absence of the IGP. **(D)** Similar output as C, with a fixed *f_cp_* value and varying α and *d_cp_*.

Over a range of predation rates (α), IGP feeding rates on the shared algal resource (f_IGP_), and death rates of the IGP (d_IGP_), we found that there is a narrow range of parameter combinations in which the R* value of the IGP can fall below that of the host snail, resulting in lower cercarial output from host snails and an observed dilution effect (Fig 4C,D). However, there is a large range of parameter value combinations in which the R* value of the IGP is above that of the host snail and disease amplification is possible, with a greater cercarial output from host snails then would be expected in the absence of the intraguild predator.

## IV. Discussion

The ecological community has the potential to mediate host-parasite dynamics via multiple mechanisms: non-host species may compete with hosts for resources, reducing host densities and thereby reducing transmission of density-dependent diseases (Keesing *et al*. 2006); parasites that seek out their host may inadvertently encounter a non-host instead (Johnson & Thieltges 2010); predators may consume hosts, parasites, or both (Borer 2002; Rohr *et al*. 2015). More than likely, multiple mechanisms that alter disease dynamics are simultaneously operating on any given host-parasite pair, making it difficult to predict how a coarse measure such as species diversity or richness will impact disease dynamics (Shaw & Civitello 2021).

In this study, we investigated the role of ecologically relevant non-host species in the snail-schistosome system. Previous attempts at using these non-host competitor snails as biocontrol in schistosome endemic regions have had mixed success (Pointier & Jourdane 2000), and there is little evidence for what may promote the success or failure of this approach (Giovanelli *et al*. 2005a). Our goal was to better understand what may be driving these mixed patterns observed in the field. We designed an experiment with two putative resource competitor species, *Melanoides tuberculata* and *Physa acuta*, and hypothesized that both would regulate host density and therefore decrease schistosome transmission potential.

*Melanoides* operated largely as we anticipated as a resource competitor. In tanks with *Melanoides*, sentinel *Biomphalaria* had a smaller body size and shed fewer cercariae. However, we generally detected weak effects of this species: *Biomphalaria* abundances and egg production were no different in tanks with *Melanoides* when compared to *Biomphalaria*-only tanks. It is possible we did not see a stronger impact of *Melanoides* snails due to a few factors: *Melanoides’* preferred habitat is sediment and while our mesocosms contained a small amount of mushroom compost as a carbon source, we were unable to add a full sediment layer because it would have inhibited exhaustive sampling of the tanks (Genner & Michel 2003). Furthermore, it is likely that our mesocosms were not able to capture the seasonal shifts of abiotic factors such as temperature and waterbody size that can result in boom-bust cycles of natural snail populations, and which may work in conjunction with interspecific competition to decrease *Biomphalaria* abundance (Rumi *et al*. 2009). However, it is also possible that our tanks more closely reflected field settings where *Melanoides* and *Biomphalaria* co-exist in the same waterbodies by occupying sediment and vegetation, respectively, such as is common in sub-Saharan Africa (Mkoji *et al*. 1992). Future work with varying abiotic conditions, habitat structural complexity, or periphyton availability may better elucidate when *Melanoides* succeeds as a biocontrol agent for *Biomphalaria* and when it fails.

*Physa acuta* is an invasive species in many regions, and has been noted to be a strong competitor with other freshwater snail species (Vinarski 2017), but similar to *Melanoides*, *Physa* has also been noted to co-exist with *Biomphalaria* in transmission sites (Laidemitt *et al*. 2022). In our experiment, *Physa* appeared to be a poor resource competitor but strong intraguild predator of *Biomphalaria* eggs. Specifically, we found that the presence of *Physa* resulted in a higher level of resources available and greater *Biomphalaria* egg production, but few new *Biomphalaria* individuals entering the population, resulting in lower abundance and an initially unexpected increase in per capita shedding of cercariae. Our generalized model of the impact of an intraguild predator on host-schistosome dynamics was congruent with what we observed in our mesocosm experiment: if a non-host snail is a weak resource competitor but can also act as an intraguild predator, then they may act as an amplifier species in the system, potentially increasing the risk of human transmission. Altogether, these mechanisms combine to create the potential to backfire as a biocontrol agent of human schistosomiasis. This finding underscores the importance of understanding the role of community members in host dynamics and parasite transmission at a mechanistic level.

Future work should focus on open questions regarding how non-host snails may influence schistosome transmission dynamics. While we were unable to infer any decoy effect of non-host snails in our experiment due to experimental issues with our tank-level parasite exposures, investigating this phenomenon would be essential to fully characterize the role that non-host snails play in this system. Furthermore, another open question we would like to highlight is the influence of non-host snails on *Biomphalaria* population size structure. *Biomphalaria* body size is tied to schistosome dynamics at multiple levels: at the time of transmission, an individual’s body size is positively correlated with parasite exposure rate and negatively correlated with susceptibility to infection given exposure (Anderson *et al*. 1982; Niemann & Lewis 1990). This phenomenon can result in interesting consequences when considering schistosome transmission within populations of differing size structures, and populations dominated by small individuals likely experience the highest rates of transmission (Shaw *et al*. 2022). However, larger snails shed greater numbers of cercariae (Civitello *et al*. 2022). In our mesocosms we found that at the individual level, *Biomphalaria* snails in tanks with *Melanoides* were significantly smaller, which could be indictive of an indirect impact of resource competition. However, tanks that contained *Physa* and *Melanoides* and *Physa* together were dominated by larger *Biomphalaria* snails producing large numbers of parasites, but these snails were exposed in laboratory conditions at identical sizes. To fully evaluate the interplay between non-hosts, host body size, and the entire schistosome life cycle, a similar mesocosm experiment that tracks population-level transmission dynamics is needed.

Despite being slated for elimination as a public health concern by the WHO by 2030, it is becoming increasingly apparent that a limited toolkit will not be sufficient to eradicate schistosomiasis (Wiegand *et al*. 2022). In order to expand our scope of interventions, it is imperative we better understand the ecology of snail hosts of schistosomes and the impact of their greater ecological community. Here, we highlight how one unexpected interaction, intraguild predation, can completely buck previous predictions about the effect of non-host snails on schistosome transmission risk to people. In addition to its relevance in our system, this finding highlights the need to better describe the types and strengths of intraspecific interactions that form the ecological backdrop of host-parasite dynamics. Moving away from coarse biodiversity metric-driven descriptions of disease dilution or amplification, and towards a goal of characterizing the community ecology of disease, has the potential to open new avenues for scientific discovery and disease mitigation.

## Author contributions

KES and DJC conceived of the question and designed the experiments. RS and RBH collected data and RS and DM processed data. KES analyzed data and wrote the manuscript with input from DJC.

## Data/scripts available

https://figshare.com/s/71603ee6b8851eed5f81

